# Malaria transmission assisted by interaction between *Plasmodium α*-tubulin-1 and *Anopheles* FREP1 protein

**DOI:** 10.1101/2019.12.16.878082

**Authors:** Genwei Zhang, Guodong Niu, Laura Perez, Xiaohong Wang, Jun Li

## Abstract

Passage of *Plasmodium* through a mosquito midgut is essential for malaria transmission. FREP1, a peritrophic matrix protein in a mosquito midgut, binds to the parasite and mediates *Plasmodium* infection in *Anopheles*. The FREP1-mediated *Plasmodium* invasion pathway is highly conserved across multiple species of *Plasmodium* and *Anopheles*. Through pulldown, nine *P. berghei* proteins were co-precipitated with FREP1-conjugated beads. After cloning these nine genes from *P. berghei* and expressing them in insect cells, six of them were confirmed to interact with recombinant FREP1 protein. Among them, *α*-tubulin-1 and heat shock protein 70 (Hsp70) were highly conserved in *Plasmodium* species with >95% identity. Thus, *P. falciparum α*-tubulin-1 and Hsp70 were cloned and expressed in *E. coli* to stimulate antibody (Ab) in mice. Our results showed that anti-serum against *P. falciparum α*-tubulin-1 significantly inhibited *P. falciparum* transmission to *An. gambiae*, while Ab against *P. falciparum* Hsp70 serum did not. The polyclonal Ab against human *α*-tubulin did not interfere formation of ookinetes, however, significantly reduced the number of *P. falciparum* oocysts in *An. gambiae* midguts. Moreover, fluorescence microscope assays showed that anti-*α*-tubulin Ab bound to impermeable *Plasmodium* ookinete apical invasive apparatus. Therefore, we propose that the interaction between *Anopheles* FREP1 protein and *Plasmodium α*-tubulin-1 directs the ookinete invasive apparatus towards midgut peritrophic matrix for the efficient passage of the parasite. *Anopheles* FREP1 and *Plasmodium* α-tubulin-1 are potential targets for blocking malaria transmission to the mosquito host.

**AUTHOR SUMMARY:** The molecular mechanisms of malaria transmission to mosquito are not well-understood. FREP1 proteins in mosquito midget PM has been proved to mediate malaria transmission by binding to parasite ookinetes. Here we reported that *Plasmodium* parasite α-tubulin-1 is an FREP1 binding partner. We initially identified the α-tubulin-1 through the FREP1-pulldown assay; Then we cloned *P. falciparum* α-tubulin-1, and demonstrated that the insect cell expressed recombinant *Plasmodium* α-tubulin-1 bound to FREP1 *in vitro*; Next, mouse anti-serum against *P. falciparum* α-tubulin-1 was found to inhibit *P. falciparum* transmission to *An. gambiae. P. falciparum* α-tubulin-1 shares >84% identical amino acid sequence with human α-tubulin, purified Ab against human α-tubulin significantly inhibited malaria transmission. Anti-human α-tubulin Ab did not interfere the gametocyte-to-ookinetes conversion. Final, we found that anti-α-tubulin Ab bound to the apical end of impermeable ookinetes. Structurally, ookinete invasive apparatus locates at the apical opening. Therefore, we propose that the interaction between *Anopheles* midgut FREP1 protein and *Plasmodium* apical *α*-tubulin-1 directs the ookinete invasive apparatus towards midgut PM for the efficient parasite invasion.

## INTRODUCTION

Recent malaria campaigns, mainly achieved by distributing insecticide-treated bed-nets, have reduced malaria cases and deaths by 30% and 47% respectively since 2000 [1]. At present, malaria control is challenged by the fast spread of insecticide-resistant mosquitoes, drug-resistant parasites [2], and the lack of malaria vaccines. New approaches for malaria control are desperately needed.

Human malaria is caused by five different *Plasmodium* species and transmitted by more than 30 different *Anopheles* species. *P. falciparum* and *P. vivax* are responsible for 99% of malaria cases. Unlike most other diseases, *Plasmodium* development in *Anopheline* mosquitoes is essential for malaria transmission. When a mosquito takes a bloodmeal from a malaria patient, haploid microgametes and macrogametes form diploid zygotes that transform into mobile ookinetes. Mobile ookinetes overcome the physical barrier of the mosquito midgut including peritrophic matrix (PM) and endothelium sequentially to initiate the infection in mosquitoes. Some interactions between a mosquito and parasites occur in the midgut, which are important for the establishment of infection. For instance, *Plasmodium vivax* Pvs25 binds to calreticulin, a mosquito midgut apical surface protein [3]. An ookinete surface enolase interacts with a midgut enolase-binding protein to mediate invasion of *P. berghei* but not *P. falciparum* [4]. So far, none of these interactions seem to be conserved across multiple species of *Plasmodium* and *Anopheles*. Discovery of a conserved pathway for malaria transmission may provide a broad spectrum target for interrupting malaria transmission through mosquitoes. Researchers have been focusing on merozoite surface proteins to develop malaria vaccines for decades [5-9]. However, vaccines based on merozoite antigens have had disappointing outcomes in clinical trials. By contrast, malaria TBV have shown promising perspectives when considering the malaria transmission bottleneck in a mosquito midgut [10].

The *FREP1* gene was discovered for its significant correlation with infection intensity of clinically circulating *P. falciparum* transmission to wild *An. gambiae* in Kenya [11]. Further investigation revealed that FREP1 bound to *P. falciparum* ookinetes to mediate their invasion [12]. Preventing *Plasmodia* from binding to FREP1 by small molecules inhibited malaria transmission [13]. Notably, Ab against FREP1 could inhibit multiple species of *Plasmodium* (e.g. *P. falciparum, P. vivax* and *P. berghei*) to invade multiple species of *Anopheles* (e.g. *An. gambiae* and *An. dirus*) [14], supporting that the FREP1-mediated *Plasmodium* transmission pathway is highly conserved across *Plasmodium* and *Anopheles* [15]. Despite the importance of FREP1-mediated *Plasmodium* invasion pathway in anopheline mosquitoes, the parasite-expressed FREP1-binding partners (FBPs) were unknown. This study aimed in identifying FBPs as a step toward elucidating the molecular pathway.

## RESULTS

### FREP1 binds to *P. berghei* parasites

Anti-FREP1 Ab prevented *P. berghei* from infecting *An. gambiae* [14]. FREP1 was able to bind to *P. falciparum* [12]. Here, we determined the interaction between FREP1 and *P. berghei*. FREP1 protein was expressed in High Five insect cells and purified using Ni-NTA column as we reported previously [12]. *P. berghei*-infected mouse blood cell lysate was used to coat microplates for enzyme-linked immunosorbent assays (ELISA), with uninfected mouse blood cell lysate was used as a control. After incubation with FREP1, anti-FREP1 Ab, and 2^nd^ Ab-conjugated with alkaline phosphatase sequentially, the enzyme activity was determined. Results showed that the absorbance changed at 405 nm (A_405_) from active FREP1 binding to infected mouse blood lysate was significantly higher than that of the negative control (*p*<0.01) (**Fig 1A**). When active FREP1 was substituted with BSA or heat-inactivated FREP1, the enzyme activity became significantly lower (**Fig 1B**), suggesting that the specific binding of active FREP1 protein to *P. berghei* was abolished. These data support that FREP1 protein binds *P. berghei* specifically.

**Figure 1.**
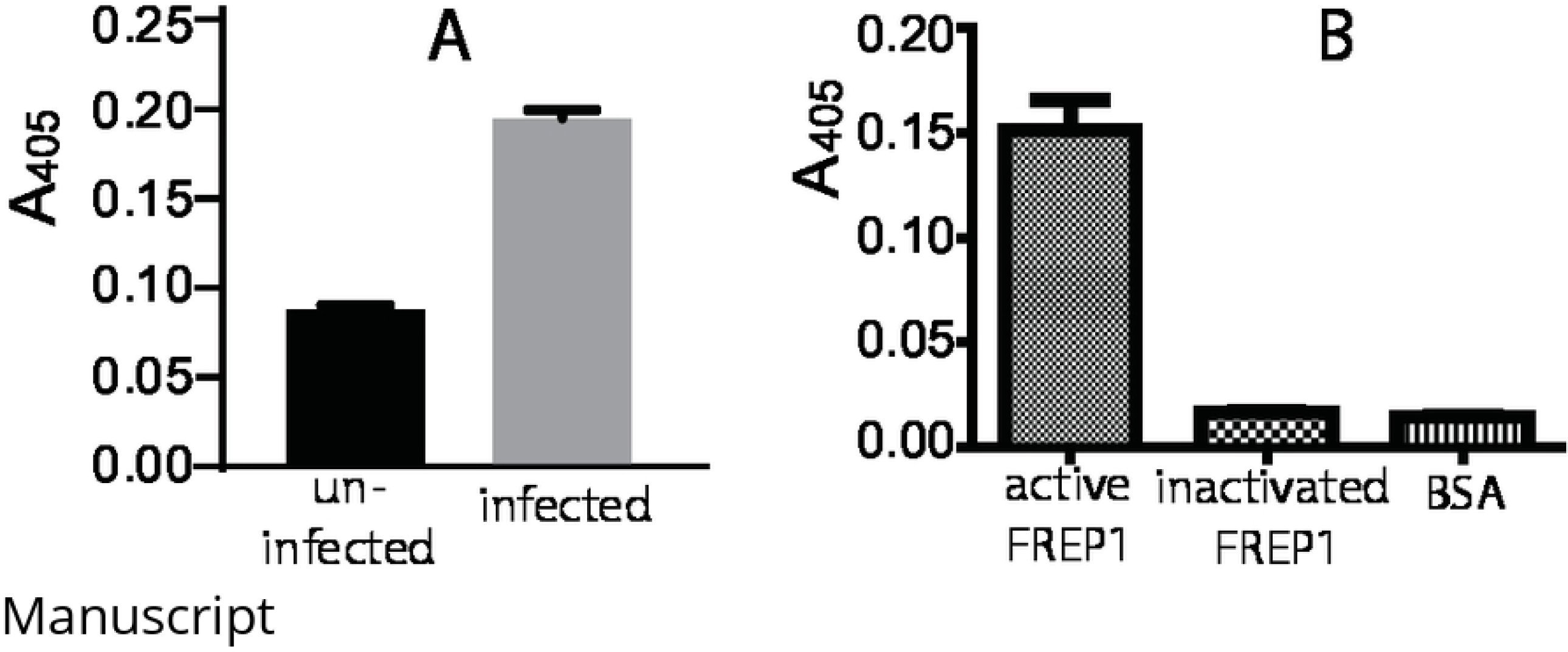
FREP1 protein bound to *P. berghei* parasites. (A) FREP1 binds to *P. berghei* infected cells significantly higher than uninfected lysate (*p*<0.001). (B) When the heat inactivated FREP1 protein or BSA replaced the functional FREP1, the binding between FREP1 and infected blood lysate disappeared.

### Separation of FBPs from *P. berghei*

To isolate FBPs, the insect cell-expressed recombinant FREP1 protein was purified to near homogeneity on a Ni-NTA column (**Fig 2A**) and covalently coupled to magnetic beads through N-hydroxysuccinimide (NHS) ester-activated chemical groups. After coupling, no detectable FREP1 was eluted from the magnetic beads (**Fig 2B**). About 13 μg of FREP1 protein was covalently linked to 1.0 mg beads. *P. berghei*-infected mouse blood lysate was incubated with the FREP1-bound magnetic beads. Magnetic beads without covalently bound FREP1, but deactivated with glycine, were used as a negative control. After washing, the bound proteins and FREP1 were eluted with reducing SDS-PAGE sample buffer and loaded onto SDS-PAGE. The result showed that some other proteins besides FREP1 appeared in the experimental lane but not in the control lane (**Fig 2C**), suggesting that FREP1 directly or indirectly bound to several proteins.

**Figure 2.**
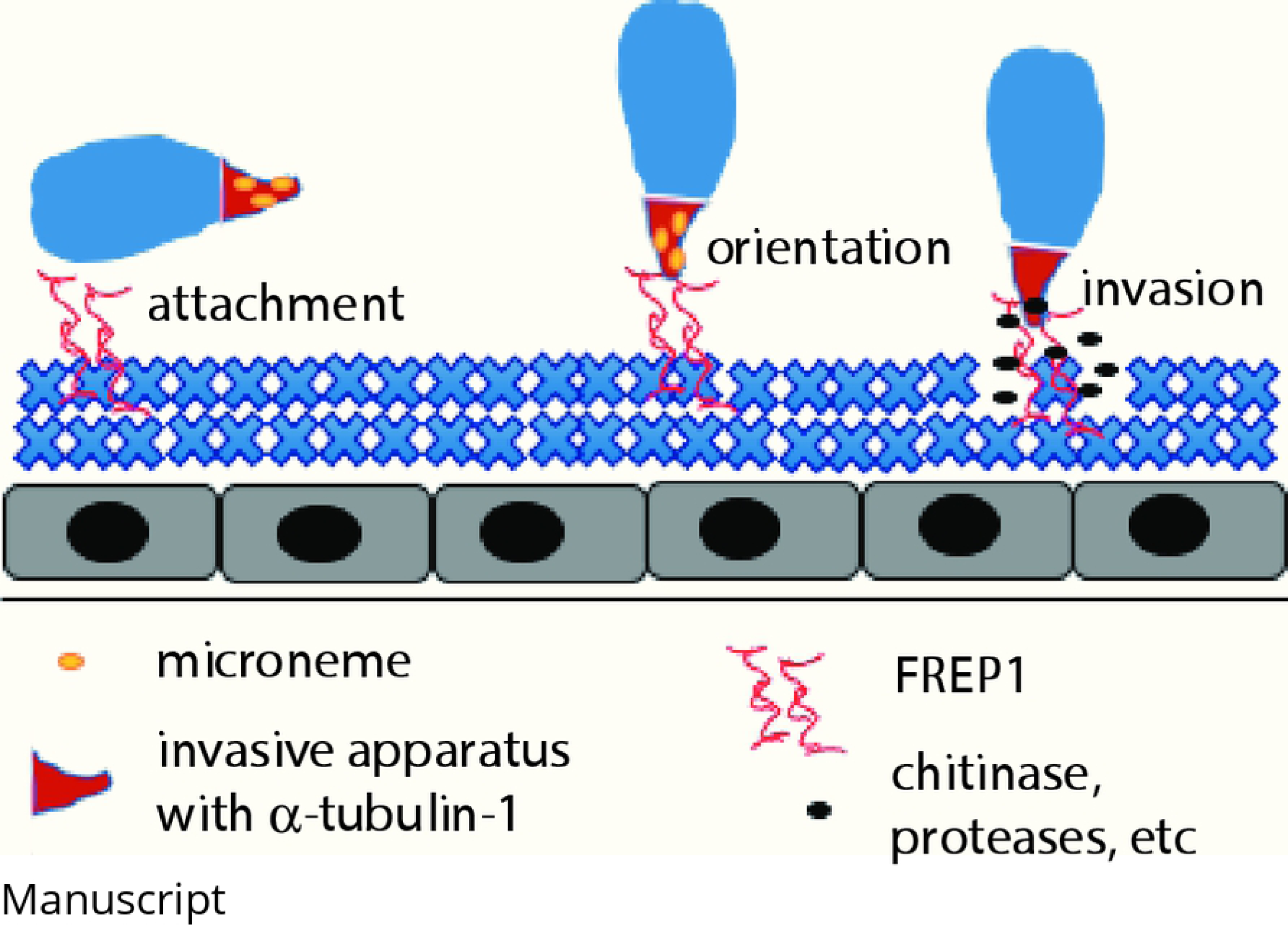
Identifying FBPs from *P. berghei.* **A**) The insect cell-expressed FREP1 protein was purified by Ni-NTA column. M: Protein molecular marker. The proteins were resolved on 12% SDS-PAGE following by Coomassie Brilliant Blue R-250. **B**) FREP1 was immobilized covalently onto the magnetic support. “Before” and “After” represent the FREP1 supernatant before coupling and after coupling with magnetic beads, respectively. **C**) Pull-down of *P. berghei-*infected cell lysate using FREP1-linked magnetic beads. The proteins were resolved on 12% SDS-PAGE following by silver staining. Lanes: 1, Pull-down of *P. berghei-*infected cell lysate by beads without FREP1 conjugated (negative control); 2, Pull-down of *P. berghei-*infected cell lysate with FREP1-conjugated magnetic beads; Marker, Protein markers; FREP1, purified FREP1 protein used to locate the recombinant FREP1 in lane 2. The rectangle highlights specific protein bands in the experimental group. This area was excised and analyzed by mass spectrometry. **D**) Determination of the interaction between recombinant FBP proteins and purified FREP1 protein by ELISA assays. The same amount of FBP1, FBP2, FBP3, FBP4, FBP5, FBP6, FBP7, FBP8, FBP9 recombinant proteins were used to coat ELISA wells. **N**: Negative control coated with insect cell-expressed CAT protein. **P**: positive control coated with *P. berghei* infected blood cell lysate. Each sample was conducted in three wells in one plate and the same experiment was conducted three times.

The gel pieces containing these different bands (black squares in **Fig 2C**) were excised from both experimental and control lanes, and analyzed by quantitative mass spectrometry. Mass-to-charge ratio of the ions was searched against the *P. berghei* (ANKA strain) protein database. Nine proteins with >99.80% identification probability (**Table 1**) were uniquely presented in the experimental samples but not in the control.

**Table 1:** Candidate FBP proteins from *P. berghei* through pull-down assays.

| **FBP Index** | **Accession # (PBANKA_)** | **Gene Name** | **MW (kDa)** | **Identification probability** |
| --- | --- | --- | --- | --- |
| **1** | 0711900 | Hsp70 | 75 | 100.00% |
| **2** | 1309700 | RNA helicase DDX5 | 59 | 100.00% |
| **3** | 0417700 | α-tubulin-1 | 50 | 100.00% |
| **4** | 1133300 | EF-1 $\alpha$ | 49 | 100.00% |
| **5** | 1316400 | U1 snRNA associated protein | 38 | 100.00% |
| **6** | 1365500 | Exported protein IBIS 1 | 37 | 100.00% |
| **7** | 0307800 | Unknown | 32 | 100.00% |
| **8** | 0701600 | Unknown | 26 | 100.00% |
| **9** | 1234500 | Unknown | 23 | 99.80% |

The coding sequences (CDS) of the nine FBP candidates were amplified by PCR from *P. berghei*, inserted into baculovirus, and expressed in High Five insect cells. The expressed recombinant proteins were used to determine their specific interaction with FREP1 by ELISA. The recombinant FBPs was coated on the ELISA plates, followed by incubation with insect cell-expressed recombinant FREP1. The retained FREP1 was detected using purified anti-FREP1 Ab. The chloramphenicol acetyltransferase (CAT) expressed in the same system was coated plate wells as the negative control. *P. berghei*-infected blood cell lysates was used to coat ELISA wells as the positive control. Results showed that six of nine FBP candidates e.g. FBP1, FBP3, FBP5, FBP6, FBP8, and FBP9 retained significantly more amount of FREP1 than that of the negative control (*p*<0.01) (**Fig 2D**), confirming they were able to bind to FREP1 protein.

### Determination of effects of candidate FBPs on malaria transmission using Ab-blocking assays

Our previous work [14] has shown that FREP1-mediated *Plasmodium* invasion pathway is conserved across multiple *Plasmodium* species including *P. falciparum, P. vivax*, and *P. berghei*. Therefore, an FBP related to malaria transmission should be conserved across the three *Plasmodium* species. We examined the sequences of the six FBPs among *Plasmodium* species. The results showed that FBP1 (Hsp70) and FBP3 (*α*-tubulin-1) were highly conserved (**Table 2**) with >98% identity across *Plasmodium* species (**File S1)**, and the conservation of other four FBP candidates was lower (<83% identity). Therefore, Hsp70 and *α*-tubulin-1 were chosen for further analysis about their effects on malaria transmission.

**Table 2:**
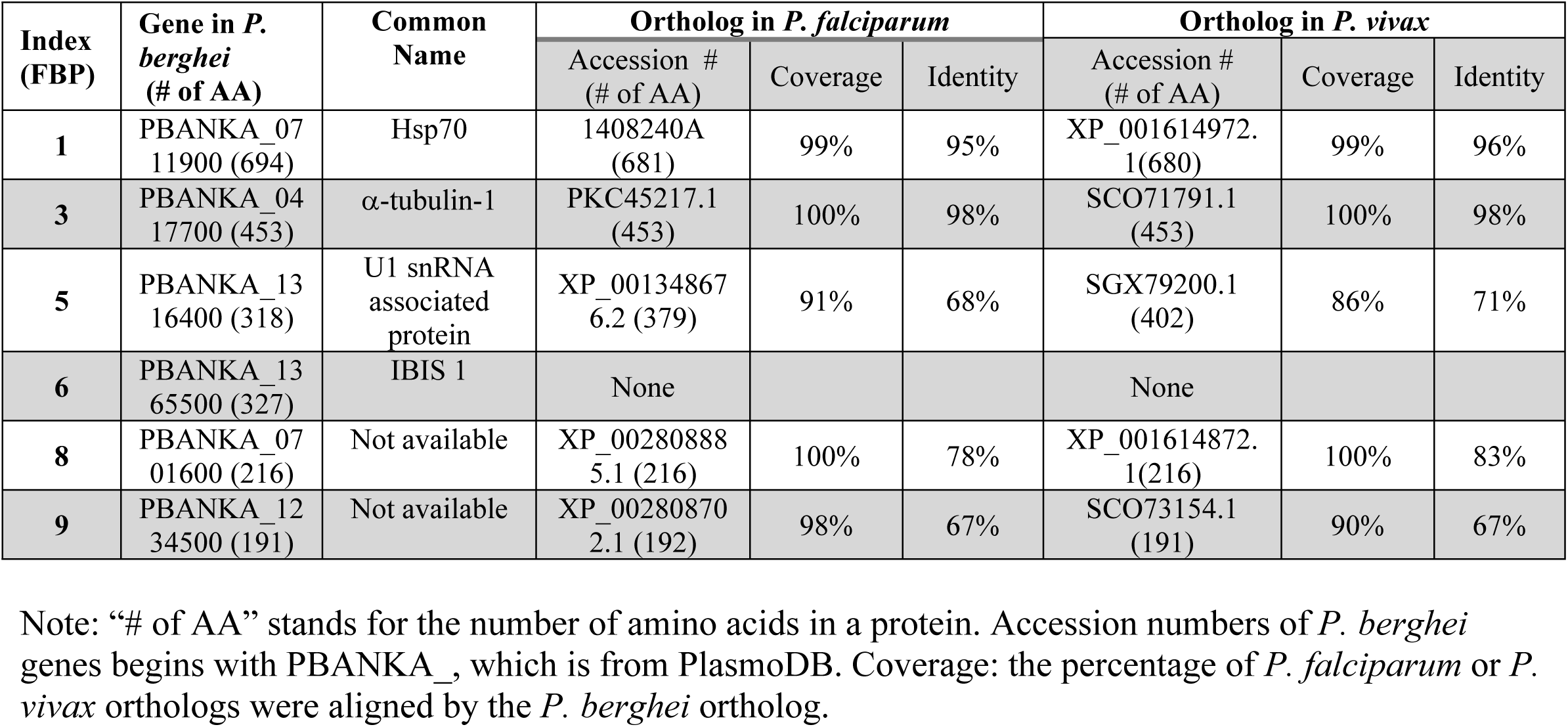
The coverage and identity of FBP orthologs among *P. berghei, P. falciparum* and *P. vivax*.

| Index (FBP) | Gene in *P. berghei* (# of AA) | Common Name | Ortholog in *P. falciparum* |  |  | Ortholog in *P. vivax* |  |  |
| --- | --- | --- | --- | --- | --- | --- | --- | --- |
|  |  |  | Accession # (# of AA) | Coverage | Identity | Accession # (# of AA) | Coverage | Identity |
| 1 | PBANKA_0711900 (694) | Hsp70 | 1408240A (681) | 99% | 95% | XP_001614972.1(680) | 99% | 96% |
| 3 | PBANKA_0417700 (453) | α-tubulin-1 | PKC45217.1 (453) | 100% | 98% | SCO71791.1 (453) | 100% | 98% |
| 5 | PBANKA_1316400 (318) | U1 snRNA associated protein | XP_001348676.2 (379) | 91% | 68% | SGX79200.1 (402) | 86% | 71% |
| 6 | PBANKA_1365500 (327) | IBIS 1 | None |  |  | None |  |  |
| 8 | PBANKA_0701600 (216) | Not available | XP_002808885.1 (216) | 100% | 78% | XP_001614872.1(216) | 100% | 83% |
| 9 | PBANKA_1234500 (191) | Not available | XP_002808702.1 (192) | 98% | 67% | SCO73154.1 (191) | 90% | 67% |

Note: “# of AA” stands for the number of amino acids in a protein. Accession numbers of *P. berghei* genes begins with PBANKA\_, which is from PlasmoDB. Coverage: the percentage of *P. falciparum* or *P.* *vivax* orthologs were aligned by the *P. berghei* ortholog.

Ab blocking assays were used to examine the effects of Hsp70 and *α*-tubulin-1 on *P. falciparum* transmission to *An. gambiae*. In order to produce sufficient proteins, we PCR-cloned *Hsp70* and *α-tubulin-1* from *P. falciparum*, expressed them in *E. coli*, and purified them with Ni-NTA columns (**Fig 3A, B**). The purified proteins were used to immunize mice. After two boosts, antisera were collected. Ab titers for Hsp70 and *α*-tubulin-1 determined by ELISA were approximately 3×10^6^ (**Fig 3C and D**). The standard membrane feeding assays were conducted using cultured *P. falciparum* mixed with 10% of antiserum. Serum from pre-immune mice served as a control. Results showed that Ab against α-tubulin-1 significantly (p<0.0004) reduced the number of *P. falciparum* oocysts in mosquitoes compared to the control (**Fig 3F**), while no inhibition was detected for anti-Hsp70 (**Fig 3E**). The experiment was repeated twice and the results were reproducible.

**Figure 3.**
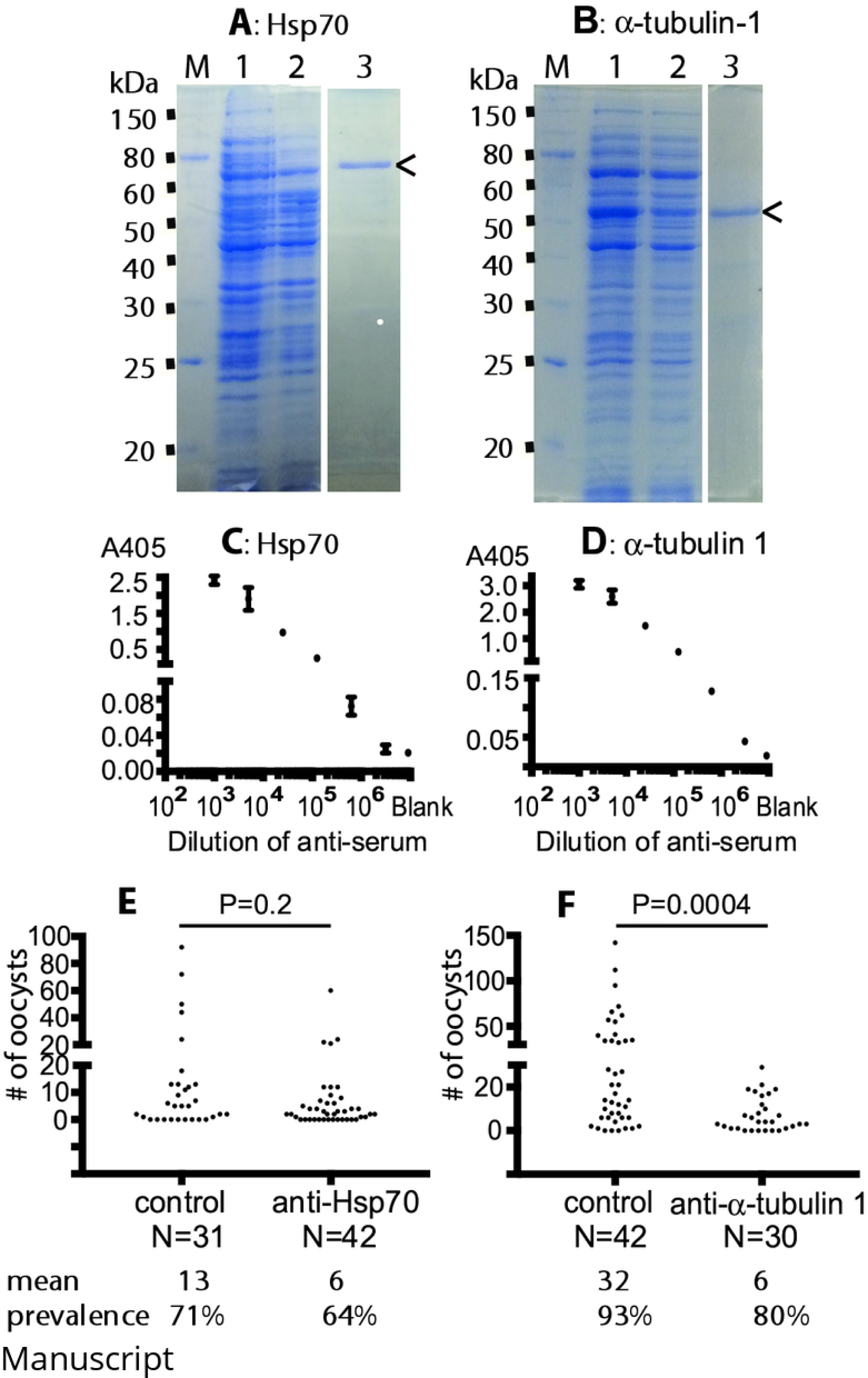
Anti-serum against *P. falciparum* α-tubulin-1 and Hsp70, and *P. falciparum* transmission-blocking assays by anti-sera. **A, B)** Expression and purification of *P. falciparum* Hsp70 and α-tubulin-1 in *E. coli.* Lanes, M: protein marker; 1: Proteins expressed after induction by IPTG; 2: *P*roteins before induction; 3: Purified *E. coli*-expressed Hsp70 or α-tubulin-1 by using Ni-NTA column, which were analyzed in a separated SDS-PAGE. **C, D**) Anti-Hsp70 and anti-*α*-tubulin-1 Ab were raised in mice respectively with two boosts. The Ab endpoint titers were measured with ELISA. X-axis coordinates show dilutions of antiserum. **E**): Anti-Hsp70 serum did not significantly inhibit *P. falciparum* transmission to mosquitoes (*p*=0.2). Each dot represents one mosquito. **F**): Mouse serum against α-tubulin-1 inhibited *P. falciparum* transmission to mosquitoes significantly (*p*=0.0004). The experiment was repeated twice and the results were reproducible.

To remove the possible effects of other components in serum, the purified polyclonal Ab was used to repeat the transmission blocking assays. However, the amount of our custom-made mouse anti-serum was very limited. Since *P. falciparum* α-tubulin-1 and human α-tubulin sequences are homologous (**Fig S1**) and they share >84% of identical amino acids (**Fig 4A**), we used the commercially available purified rabbit polyclonal Ab against human *α*-tubulin in the Ab transmission-blocking assays. The purified Ab was mixed with *P. falciparum*-infectious blood and fed to *An. gambiae*. A non-related purified rabbit polyclonal Ab (anti-V5 tag) was used as a control. At the end, the number of oocysts in mosquito midguts was counted. Results showed that Ab against human α-tubulin significantly reduced the number of *P. falciparum* oocysts in *An. gambiae* midguts (*p*<0.015) compared to the negative control (**Fig 4B**). Notably, the infection prevalence rate was decreased from 27% and 68% to 6% and 4% respectively in two replicates. In fact, only one or two mosquitoes had one oocyst in experimental groups.

Ab targeting β-tubulin were also used in the transmission-blocking assays. Even though human α-tubulin and human β-tubulin share 41% identical amino acid sequences (**Fig 4A**), there was no significant difference of oocyst counts between experimental mosquito midguts and the control (p>0.23), indicating that the polyclonal Ab against human β-tubulin did not prevent *P. falciparum* from invading *An. gambiae* (**Fig 4B**).

**Figure 4:**
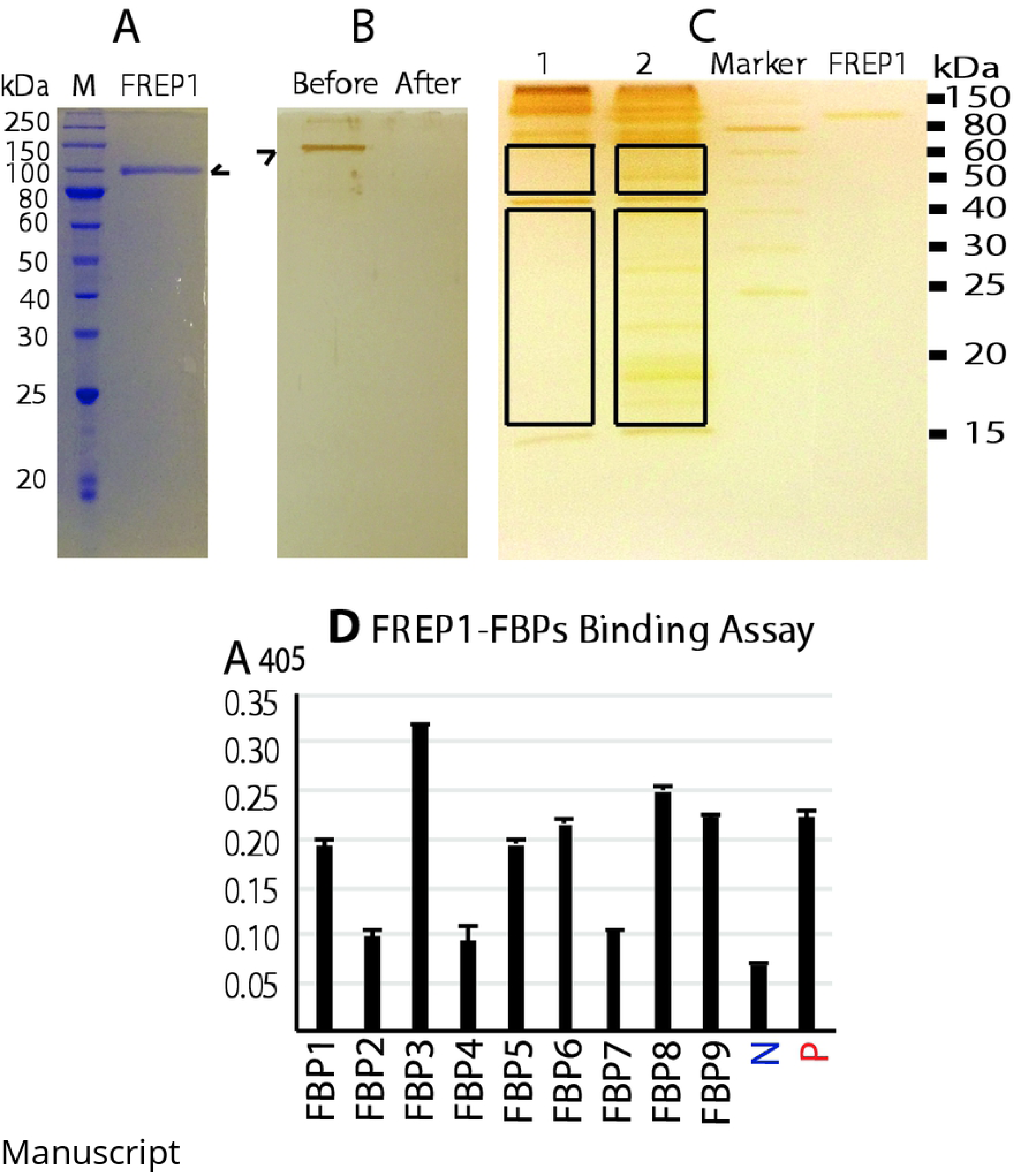
Protein sequence comparison and purified Ab transmission-blocking assays. **A**) Sequence alignment of human α-tubulin (HA), *P. falciparum* α-tubulin-1 (PfA), human β-tubulin (HB), *P. falciparum* β-tubulin (PfB). **B**) Purified rabbit anti-human α-tubulin polyclonal Ab (labeled with α) significantly reduced the number of *P. falciparum* oocysts in *An. gambiae* midguts, while purified Ab against β-tubulin (labeled with β) did not inhibit *P. falciparum* transmission to *An. gambiae.* A non-related purified rabbit polyclonal Ab (anti-V5, labeled with control) was used as the negative control. Wilcoxon test was used to calculated P-value. The experiment was repeated twice and the results were similar.

### Anti-α-tubulin Ab did not affect the conversion of *P. falciparum* gametocytes to ookinetes

There is a possibility that the reducing number of oocysts in mosquitoes by Ab was caused by the lower gametocyte-to-ookinete conversion. To determine this, the purified rabbit polyclonal Ab against human α-tubulin was added into the *P. falciparum* culture containing ∼3% gametocytes. The same amount of non-related purified rabbit Ab (anti-V5) was used as a control. After incubation, the ookinetes stained with the Giemsa staining were examined. The ookinetes were found in both the experimental group (**Fig 5A**) and the control group (**Fig 5B**) under the microscopy bright field. The ookinete conversion rates (CR), e.g. the percentage of ookinetes among the total gametocytes, were calculated. The ookinete CR with anti-α-tubulin Ab was 12.8%, and the ookinete CR with anti-V5 Ab was 12.2% (**Table 3**). The data showed that Ab against human α-tubulin did not significantly affect *P. falciparum* gametocytes to ookinetes comparing to that of the negative control (p>0.1).

**Table 3:**
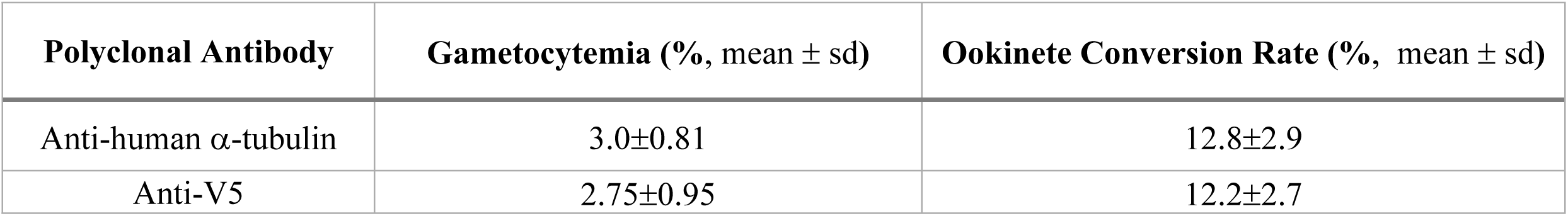
The effects of antibodies on ookinete conversion.

**Figure 5:**
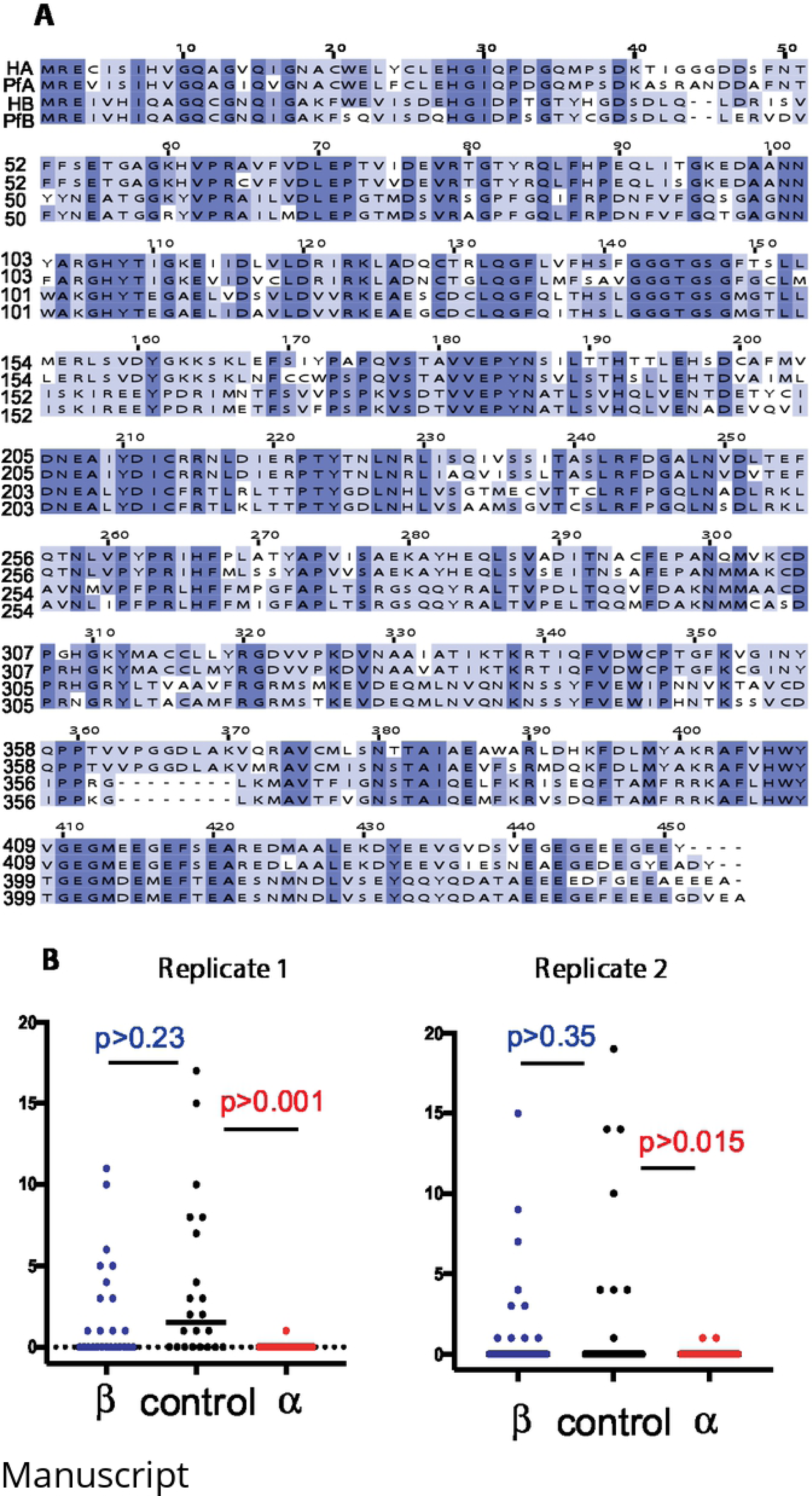
The production *P. falciparum* ookinetes in the present of different antibodies.

## ANTI-α-TUBULIN AB BOUND TO OOKINETE APICAL INVASION APPARATUS

The Ab transmission-blocking assays strongly supported that ookinete α-tubulin-1 was exposed on the cell surface. Determination of the localization of α-tubulin-1 on impermeable ookinetes would further elucidate the molecular mechanisms of this pathway.

We labelled polyclonal Ab against human α-tubulin with CF™ 568 dye, and followed the nonpermeable approach for immunofluorescence assays (IFA) [12] to investigate the interaction between Ab and ookinetes. *P. falciparum* ookinetes were cultured *in vitro,* deposited onto slide coverslips, and fixed with paraformaldehyde to acquire impermeable cells. Then these cells were incubated with fluorescence labelled anti-human α-tubulin Ab. Finally, the cells were examined under a fluoresces microscope. Results showed than anti-α-tubulin Ab bound to the apical end of ookinetes (pointed by white arrows) (**Fig 6B, Row 1**), but not the gametocyte (pointed by the green arrow) (**Fig 6B, Row 1)**. The ookinetes and gametocyte were confirmed by DAPI staining (**Fig 6B, Row 2**) and bright field (**Fig 6B, Row 3**). The DAPI also stained many free parasites and asexual stage parasites. An unrelated Ab, anti-V5 polyclonal Ab was used to replace anti-human-α-tubulin polyclonal Ab as negative control. As expected, anti-V5 Ab did not stain any ookinete (**Fig 6A**).

**Figure 6:**
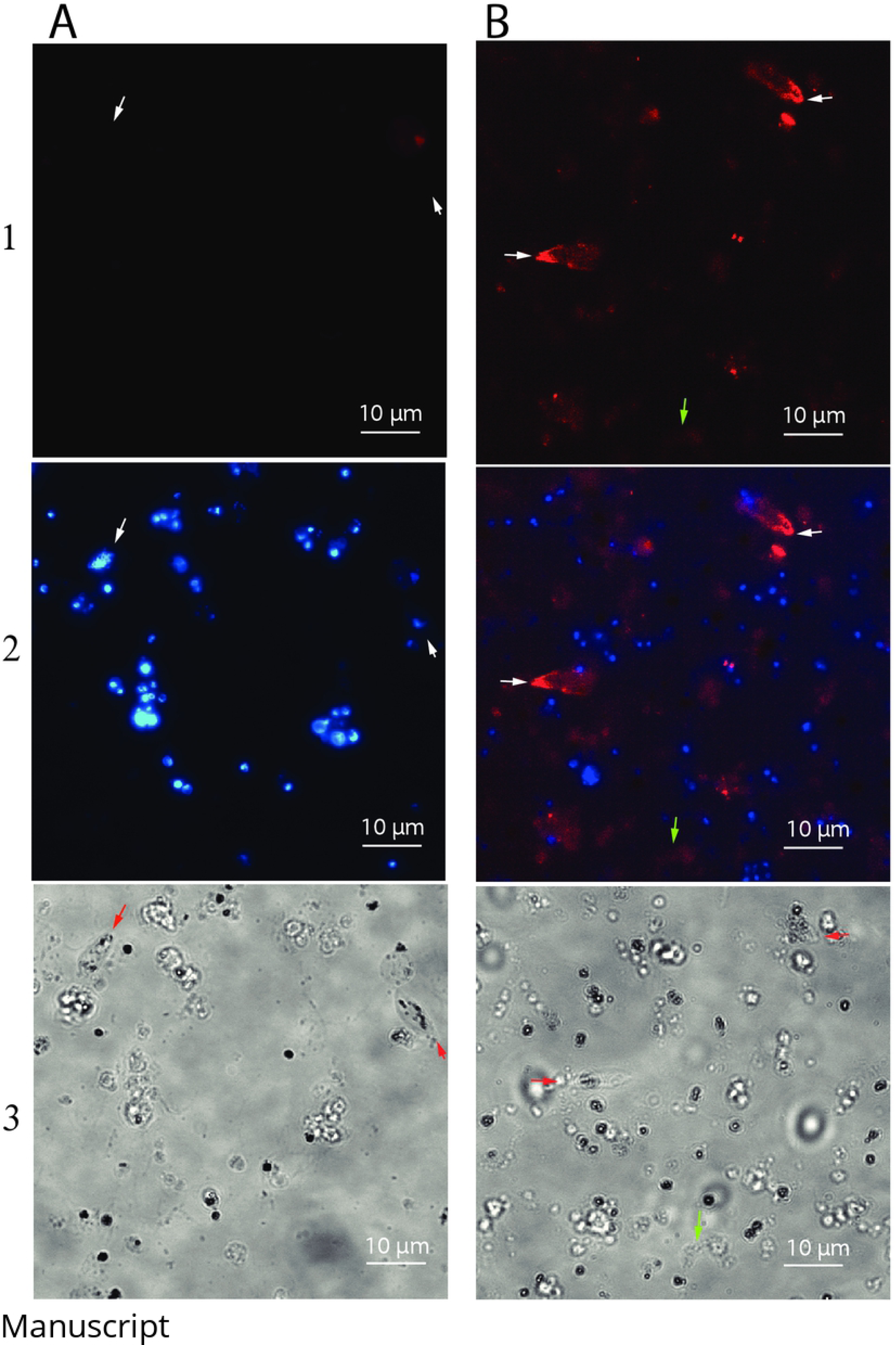
Immunofluorescence assay showed that anti-human α-tubulin Ab bound to the impermeable-fixed *P. falciparum* ookinetes at their apical end. **A**) Negative control. *P. falciparum* ookinetes stained by an un-related polyclonal Ab (ant-V5 AB). No binding signals were detected. **B**) Impermeable ookinetes stained by the anti-human-α-tubulin Ab. Ab bound to the apical end of a paraformaldehyde-fixed ookinete. In each treatment, the 1^st^ row detected α-tubulin-1 (red color). The 2^nd^ showed the co-localization of *P. falciparum* (nuclei, blue color) and α-tubulin-1. The 3^rd^ column shows the bright views of the cells. The white arrows point to ookinetes. The green arrow points to a gametocyte.

## DISCUSSION

Although important for malaria control, molecular mechanisms related to malaria transmission to mosquitoes have rarely been reported. So far, only the FREP1-mediated *Plasmodium* transmission pathway has been proven to be conserved across multiple *Plasmodium* species transmission to multiple *Anopheles* mosquitoes [15]. This conservation is confirmed by transgenic *An. gambiae*. Knocking out FREP1 gene renders *An. gambiae* resistant to multiple species of *Plasmodia* [16]. However, the molecular mechanism of FREP1-mediated *Plasmodium* invasion pathway is incomplete if parasite-expressed FBP is unknown. This study aimed to identify the FBP.

Identifying an unknown binding partner or receptor is always a challenge [17] and the discovery of FBPs is further complicated by the limited source of materials. Less than 5% of *Plasmodium*-infected cells eventually develop into gametocytes, less than 1% of gametocytes become ookinetes [10]. Moreover, a method to separate ookinetes from other cells has not been developed yet [18]. A large amount of non-specific proteins presented in original materials for pull-down assays are expected. We tried many approaches. Eventually magnetic beads conjugated with the purified insect cell-expressed FREP1 reduce pulldown noise. Searching differential mass spectrometry signals against *P. berghei* protein database further eliminated more non-related proteins. Finally, the Multidimensional Protein Identification approach followed by statistical evaluation [19] provided a reliable short list for further analysis. We cloned and expressed the candidate genes in insect cells, and recombinant proteins were used to confirm the interactions between FREP1 and FBP candidates. Interestingly, six recombinant FBP candidates bound to FREP1 protein. Among them, Hsp70 and α-tubulin-1 are extremely conserved among *Plasmodium* species, consistent to the conservation of FREP1-mediated *Plasmodium* transmission pathway. Thus, Hsp70 and α-tubulin-1 were selected to determine their function on *Plasmodium* transmission.

Through transmission-blocking assays, we found that customized antiserum against *P. falciparum* α-tubulin-1 as well as purified polyclonal Ab significantly inhibited *P. falciparum* transmission to *An. gambiae,* while Ab against Hsp70 did not. Moreover, purified rabbit Ab against α-tubulin reduced *P. falciparum* oocysts nearly to zero, while anti-β-tubulin Ab did not prevent *P. falciparum* transmission. It is no doubt that the interaction between *Anopheles* FREP1 and *Plasmodium* α-tubulin-1 involves in malaria transmission. Next, we examined whether anti-α-tubulin Ab interfered with ookinete production, and found that anti-α-tubulin Ab did not interfere with the gametocyte conversion to ookinetes. Therefore, anti-α-tubulin Ab in serum shall have access to the living ookinetes and inhibited *Plasmodium* ookinete to infect mosquito midguts.

To further elucidate the molecular mechanism of FREP1-α-tubulin-1 interaction facilitating parasite transmission, IFA was used to view the interaction between Ab and ookinete α-tubulin-1. The results demonstrated that anti-α-tubulin Ab bound to the apical end of impermeable ookinetes. The ookinete apical end is the invasive apparatus [20, 21].

Collectively, we propose that an ookinete uses the following steps to overcome the physical barrier of PM. First, *Anopheles*-expressed FREP1 protein at the midgut PM binds to *Plasmodium* ookinete-expressed molecules. This action anchors ookinetes on PM. Then, the interaction between FREP1 and *α*-tubulin-1 at the apical invasion apparatus directs the ookinete invasive apparatus toward the midgut PM. This action increases the efficiency of the secreted digestive enzymes [21] to disrupt the integrity of the PM (**Fig 7**). Final, the parasite penetrates the PM and invades midguts.

**Figure 7:**
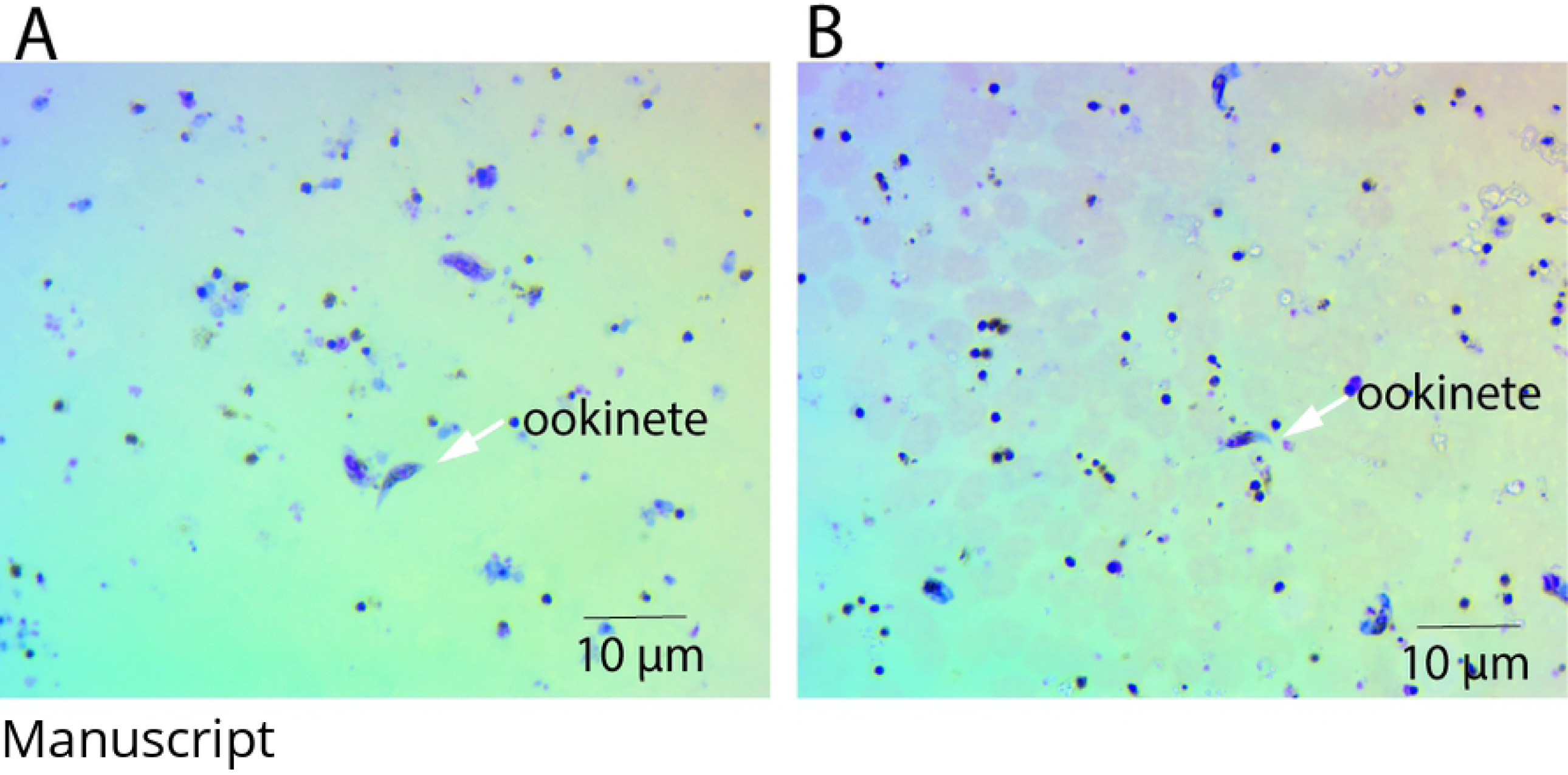
A hypothetical model of FREP1-α-tubulin-1 interaction that mediates the *Plasmodium* transmission to *Anopheles*. From left to right: initial attachment of parasites on mosquito midgut peritrophic matrix, mediated by several types of interactions, anchors ookinetes to PM; orientation of ookinete invasion apparatus, mediated by FREP1-α-tubulin-1 interaction, directs invasive apparatus opening toward PM; and disruption of PM integrity, caused by the enzymes released from the parasitic apical opening, facilitates a parasite for invasion.

In summary, we elucidate a FREP1-mediated *Plasmodium* invasion pathway in *Anopheles* midguts and discovers *Plasmodium* α-tubulin-1 as a FREP1 binding partner. Direct interaction between FREP1 and α-tubulin-1 at ookinete invasive apparatus is hypothesized to orientate the parasite for efficient invasion. This discovery provides targets for malaria control, particular vaccines and drugs that block malaria transmission.

## MATERIALS AND METHODS

### Ethics Statement

This study was carried out in strict accordance with the recommendations in the Guide for the Care and Use of Laboratory Animals of the National Institutes of Health. The protocol was approved by the Animal Care and Use Committee of the University of Oklahoma (permit number R15-012) and the Animal Care and Use Committee of Florida International University (permit number IACUC-16-073).

### Determining the interaction between insect cell-expressed recombinant FREP1 and *P. berghei* by ELISA

High Five insect cell-expressed FREP1 and anti-FREP1 polyclonal Ab was obtained as we previously reported [12]. GFP transgenic *P. berghei* (ANKA strain) was obtained from the American Type Culture Collection (ATCC, Manassas, VA). Swiss Webster mice were purchased from Envigo (Indianapolis, IN). Mice were infected with *P. berghei* through i.p. injection. The parasitemia was checked every other day using Giemsa staining of thin blood smears. When the parasitemia reached 10%, gametocytes were induced by i.p. injecting mice with 60 mg phenylhydrazine hydrochloride (Santa Cruz Biotechnology, Dallas, Texas) per kilogram body weight (4mg/mL, dissolved in PBS). Two days later, *P. berghei*-infected red blood cells and uninfected red blood cells were collected, washed three times with PBS, and re-suspended in PBST (1xPBS containing 0.2% Tween-20). The lysates were prepared by ultra-sonication of cells 6 times with 10 seconds (s) each and 50 s pause on ice, followed by centrifugation at 8,000 ×g for 5 minutes (m) to remove intact cells and insoluble aggregates. The proteins in supernatants were used for ELISA assays and pulldown assay.

The ELISA method used to study FREP1-parasite interaction has been described in details before [12]. A 96-well plate (Brand, Wertheim, Germany) was coated with 2 mg/mL lysates (measured by Bradford assay) and incubated overnight at 4°C. The next day, each well was incubated with the following solutions: 200 µL blocking buffer (2 mg/mL BSA in PBS) for 1.5 hour (hr), 100 µL recombinant FREP1 protein (100 µg/mL) at room temperature (RT) for 1 hr, 100 µL of purified anti-FREP1 Ab (1:2,000 dilution with PBST) for 1 hr at RT, and 100 µL of alkaline phosphatase-conjugated anti-rabbit IgG (1:20,000 dilution in PBST) for 45 m at RT. The wells were washed with PBST three times between incubations. In the end, the plates were developed with 100 µL of *p*-NPP solution (Sigma-Aldrich, St. Luis, MO) until the colors appeared. Finally, A_405_ was measured. Each experiment was repeated three times and the t-test was used to compare the difference between the control and the experimental group.

### Immobilizing FREP1 onto magnetic beads

About 300 µL of N-hydroxysuccinimide (NHS) ester-activated magnetic beads (Thermo Fisher, Weston, FL) were placed into a 1.5 mL microcentrifuge tube on a magnetic stand, and the supernatant was discarded. Beads were then washed with 1 mL of ice-cold 1 mM hydrochloric acid with gentle vortex for 15 s. After discarding the supernatant, 300 µL of purified FREP1 protein (in 50 mM borate, pH 8.5) was added immediately and vortexed for 30 s. The mixture was incubated for 1-2 hr at RT on a rotator. During the first 30 m of the incubation, the tube was vortexed for 15 s every 5 m. For the remaining time, the tube was vortexed for 15 s every 15 m. FREP1-coupled magnetic beads were then washed twice with 1 mL of 0.1 M glycine (pH 2.0) for 15 s and once with 1 mL of ultrapure water for 15 s. Next, 1 mL of Quenching Buffer (3M ethanolamine, pH 9.0) was added, vortexed for 30 s, and incubated for 2 hr at RT on a rotator. Finally, FREP1-coupled magnetic beads were stored in 300µL Storage Buffer (50 mM borate with 0.05% sodium azide) at 4°C. The coupling efficiency assessment was performed using Bradford assay measuring the amount of FREP1 proteins in the solution before and after coupling.

### Affinity chromatography and quantitative mass spectrometry to identify FBP proteins

About 1 mL *P. berghei*-infected cell lysate was incubated with 50 µL FREP1-coupled magnetic beads at 4°C for 2 hr with gentle rotation. The beads were then washed three times to remove non-specifically bound proteins with Pierce IP Lysis Buffer (catalog number: 87787). The retained proteins were eluted with 50 µL SDS-PAGE loading buffer. In the control group, procedures were the same as the experimental group, with the exception of the use of glycine-deactivated NHS ester-magnetic beads to replace FREP1-coupled beads. The elusion samples were analyzed with 12% SDS-PAGE. The SDS-PAGE gel was stained with the Coomassie-brilliant blue R250. Specific protein bands in SDS-PAGE gel were excised and analyzed by quantitative mass-spectrometry (Oklahoma State University Mass Spectrometry Service Center). The signals were searched against *P. berghei* protein database (plasmodb.org) to identify peptide hits. The difference in protein abundance between the control and the experiment group was obtained using a published approach [19]. Briefly, the spectral count data were collected using Multidimensional Protein Identification Technology. The Fisher Exact Test is used to calculate the difference p-value using Scaffold (Proteome Software Inc). The calculated p values were transformed to probabilities by an equation of probability equals to (1-*p*)x100.

### Expression of active candidate proteins using baculovirus in insect cells to verify protein-protein interaction

The coding sequences (CDS) of the nine FBP candidates were amplified by PCR from *P. berghei* using gene-specific primers (**Table 4**). Depending on primer sequences, PCR products were digested with restriction enzymes Bam H1 and Xba I (New England BioLabs, MA) or Bam HI and Xho I (New England BioLabs, MA). The column isolated DNA fragments were ligated into our modified pFastBac1 plasmid having 6xHis tag at C-terminus, and positive recombinant plasmids were transformed into DH10Bac competent cells. Recombinant bacmids were extracted from white colonies. Following the manufacturer’s manual [22], 0.5 mL 2×10^5^ /mL sf9 cells in complete medium (Grace’s medium (ThermoFisher) containing 10% FBS) were deposited into one well of a 24-well plate. After incubation for 30 m, the supernatant was removed. Meantime, one μL bacmid DNA (1 μg/μL) combined with 25 μL Grace’s medium was mixed gently, and 2μL Cellfectin II (ThermoFisher) diluted with 25 μL Grace’s medium was mixed gently. The DNA mixture and Cellfectin II mixture were combined and incubated for 30 m at RT, and then added to the above prepared sf9 cells. After incubation for 5 hr at 27 °C, the transfection mixture was removed and 0.5 mL complete growth medium (Grace’s medium with 10% FBS and 1x Penicillin/Streptomycin) was added. After incubation at 27 °C for three days, the medium containing recombinant baculovirus particles was collected and used to infect High Five insect cells to express recombinant proteins. After three passages (10 μl old cell culture to 200 μl new cell culture), expressed proteins were confirmed with ELISA by detecting the His-tag at the C-terminus of recombinant proteins.

**Table 4:**
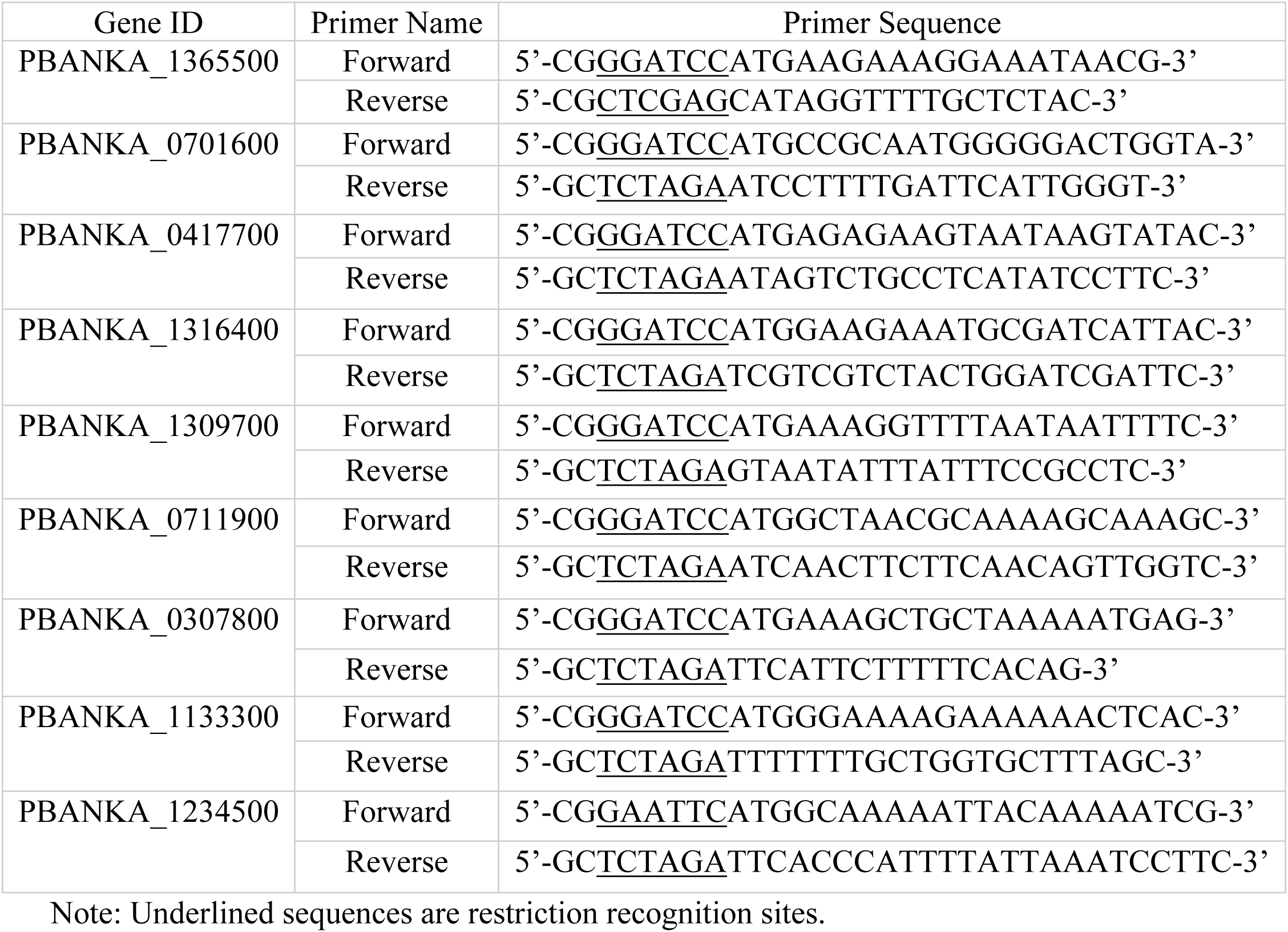
Primers for FBPs expression using baculovirus system.

### Determination of FBPs-FREP1 interaction by ELISA

The expressed recombinant FBP proteins were used to determine their specific interaction with FREP1 by ELISA. The expression levels of different FBPs in the cell lysate were normalized to 1 nM and then a 96-well plate was coated overnight with 100 μl cell lysate to detect interactions with FREP1. After blocking the plate with 100 μl 0.2% BSA in 1×PBS, 100 μl recombinant FREP1 in PBS (1 nM) was introduced onto the wells and incubated for 1.5 h with the plate at RT. Purified anti-FREP1 Ab (3 μg/mL in PBS) was used to detect any retained FREP1. Followed by 100 μl 2^nd^ Ab (1:10,000 dilution in PBS). The plates were developed with 100 µL of *p*-NPP solution (Sigma-Aldrich, St. Luis, MO) until the colors appeared. The absorbance at 405 nm was measured using a plate reader. Between each incubation, wells were washed with 100 µL PBST three times with 3 m incubation each time. For the control group, 1 nM of CAT protein (Chloramphenicol acetyltransferase, containing a 6×His tag at C-terminus) expressed in the same baculovirus-expression system (ThermoFisher) was used to coat plate wells. The *P. berghei*-infected mouse blood cell lysate (1 mg/ml proteins) was used to coat wells as the positive control. Each sample was repeated in three wells and the t-test analysis was used to difference between the negative control and experimental group.

### Production of Ab against FBPs

To generate sufficient amount of proteins for immunonization, the coding sequence (CDS) of *P. falciparum HSP70* was amplified with primers of 5’-CG<u>GGATCC</u>ATGGCTAACGCAAAAGCAAAGC-3’ and 5’-CGC<u>AAGCTT</u>TTAATCAACTTCTTCAACAGTTGGTC-3’. The CDS of *P. falciparum* α-tubulin-1 was amplified with primers of 5’-CG<u>GGATCC</u>ATGAGAGAAGTAATAAGTATAC-3’ and 5’-GC<u>TCTAGA</u>ATAGTCTGCCTCATATCCTTC-3’. The sequences underlined are restrict enzyme recognition sites. The DNA fragments were digested with corresponding restriction enzymes and ligated into the pQE30 plasmid. Recombinant proteins were expressed in *Escherichia coli* M15 strain induced by 1 mM isopropyl 1-thio-β-D-galactopyranoside (IPTG). After 3-4 hr induction at 37°C, cells were lysed in buffer B (8 M urea, 100 mM NaH_2_PO_4_, 10 mM Tris-Cl, pH 8.0). Recombinant FBPs were purified on Ni-NTA column using a standard protocol [23]. Customized polyclonal Ab was generated in Swiss Webster mice by priming with 20 μg purified recombinant FBPs in CFA adjuvant (v/v, 1:1), followed by two additional boosts (with 2-weeks intervals) with 20 μg proteins in IFA adjuvant (v/v, 1:1). An identical volume of buffer B with the same adjuvant was used as a negative control to immunize mice. Fifty days later, mouse sera were collected. Ab titers were checked using ELISA with a serial dilution of the collected sera added into the 96-well plates that had already been coated with the corresponding antigens. Equivalent amounts of control mouse serum were used as negative controls. The cut-off criterion was the dilution that OD_405_ signal decreases to the mean of negative control replicates plus 2×S.D.

### Culture *P. falciparum* in lab

*P. falciparum* parasites (NF54 strain from MR4, Manassas, VA) were maintained in 5 mL RPMI 1640 medium (Life Technologies) supplemented with 10% heat-inactivated (56 °C for 45 min) human AB+ serum (Interstate Blood Bank, Memphis, TN), 12.5g/ml hypoxanthine, and 4% hematocrit using a 6-well cell culture plate (Corning, NY) in a candle jar at 37 °C [12]. The parasitemia or gametocytemia was analyzed every day by Giemsa staining of thin blood smears. When the parasitemia reaches 5%, 0.5 ml of culture was transferred into 5ml fresh complete RPMI-1640 supplemented with 4% hematocrit. The remaining culture was maintained for another 10 days to two weeks before using it to infect mosquitoes. To prepare ookinetes, *P. falciparum* cultures harboring 2-5% stage V gametocytes were diluted 10-fold in complete RPMI 1640 without sodium bicarbonate and incubated at room temperature for 24 h as described [25].

### Ab transmission-blocking assays of *P. falciparum* infection in *An. gambiae*

Standard membrane feeding assays (SMFA) were performed as the previous reports [12, 24]. About 30 μL anti-sera were mixed with 330 μL cultured *P. falciparum* and fed 3-5-day-old female *An. gambiae* mosquitoes. After feeding, the engorged mosquitoes were maintained with 8% sugar (w/v) at 27°C. Seven days after infection, mosquitoes were dissected, and the midguts were stained with 0.2% mercurochrome for 30 min. Then, the number of oocysts was counted under light microscopy. Nonparametric statistical analysis, e.g. Wilcoxon test that was implemented in GraphPad Prism software (Version 6h, San Diego, CA) was used to analyze the inhibition effect of Ab.

To rule out the effect of other components in sera, the purified rabbit polyclonal Ab (Abcam, Cambridge, MA) was used in malaria transmission blocking assay. About 6μL rabbit polyclonal Ab against human *α*-tubulin (0.3 mg/mL) was added into 200 μL *P. falciparum*-infectious blood (final Ab concentration was about 0.01 mg/mL) and fed to *An. gambiae*. The same amount of non-related purified rabbit polyclonal Ab (anti-V5 tag) was used as a control. Seven days late, the number of oocysts in mosquito midguts was counted for statistical analysis by Wilcoxon test.

### The ookinete conversion assays by Ab

First, the number of gametocytes in cultures was counted under microscope using the blood smear. About 1 mL 15-day *P. falciparum*-cultured blood that harbors stage V gametocytes was collected by centrifugation (800 xg for 4 m), and resuspended into 10-fold ookinete cultures (complete RPMI-1640 without NaHCO_3_). About 3 µL of polyclonal anti-human α-tubulin Ab (0.3 mg/mL, Abcam) was then added into 150 µL *P. falciparum*-infectious blood (final Ab concentration was about 6 µg/mL) to a well in a 96 well plate. Then the parasites were incubated under 21-23 °C for about 16 hours. The same amount of non-related purified rabbit Ab (anti-V5) was used as a control. The number of ookinetes was counted, and the ookinete conversion rate (CR), the percentage of ookinetes among the total *P. falciparum* gametocytes, was calculated.

### Determine the localization of α-tubulin-1 on *P. falciparum* ookinetes by IFA

We labelled rabbit polyclonal Ab against human α-tubulin (ProteinTech, IL, USA) with CF™ 568 dye (Mix-n-Stain™ CF® Dye Antibody Labeling Kits - CF®568, Biotium Inc, CA), and followed the nonpermeable approach for IFA [12] to investigate the interaction between Ab and ookinetes. The cultured *P. falciparum* ookinetes were deposited onto slide coverlips, and fixed with 4% paraformaldehyde in PBS for 30 m to acquire impermeable cells. Before the cells were completely dry, these cells were blocked with 0.2% BSA in PBS for 90. Then the cells were incubated with fluorescence labelled anti-human-α-tubulin Ab (1:250 dilution in PBS containing 0.2% BSA. The final Ab concentration was 0.01mg) in the darkness at room temperature for two hrs. Between each incubation, the cells were washed 3 times with PBS containing 0.2% BSA (5 m each wash). Finally, coverslips were then rinsed in distilled water for 20 s and mounted on glass slides using 50 μL of Vectashield mounting media (Vector Laboratories, Burlingame, CA). Finally, cell staining was examined using a Nikon Eclipse Ti-S fluorescence microscope.

## AUTHOR CONTRIBUTIONS

JL conceived the concept and designed the study. GZ, GN, LP, and JL conducted experiments. XW analyzed some sequences. GZ and JL wrote the manuscript. All authors edited the manuscript.

## ACKNOWLEDGMENT

This study was supported by NIAID 1R01AI125657, 1R21AI115178, and NSF Career Award 1453287. The Mass Spectrometry Service Center at Oklahoma State University analyzed samples from SDS-PAGE.

## FIGURE LEGENDS

**Figure S1**: **The sequence alignment of human α-tubulin and *P. falciparum* α-tubulin-1**.

## REFERENCES

1. WHO. Malaria: World Health Organization; 2018 [cited 2018 May 26]. 20 April 2018:[Available from: http://www.who.int/news-room/fact-sheets/detail/malaria.

2. Yuan J, Cheng KC, Johnson RL, Huang R, Pattaradilokrat S, Liu A, et al. Chemical genomic profiling for antimalarial therapies, response signatures, and molecular targets. Science. 2011;333(6043):724–9. Epub 2011/08/06. doi: 10.1126/science.1205216. PubMed PMID: 21817045; PubMed Central PMCID: PMCPMC3396183.

3. Rodriguez Mdel C, Martinez-Barnetche J, Alvarado-Delgado A, Batista C, Argotte-Ramos RS, Hernandez-Martinez S, et al. The surface protein Pvs25 of Plasmodium vivax ookinetes interacts with calreticulin on the midgut apical surface of the malaria vector Anopheles albimanus. Molecular and biochemical parasitology. 2007;153(2):167–77. doi: 10.1016/j.molbiopara.2007.03.002. PubMed PMID: 17442413.

4. Vega-Rodriguez J, Ghosh AK, Kanzok SM, Dinglasan RR, Wang S, Bongio NJ, et al. Multiple pathways for Plasmodium ookinete invasion of the mosquito midgut. Proceedings of the National Academy of Sciences of the United States of America. 2014;111(4):E492–500. doi: 10.1073/pnas.1315517111. PubMed PMID: 24474798; PubMed Central PMCID: PMC3910608.

5. Yoshida S, Nagumo H, Yokomine T, Araki H, Suzuki A, Matsuoka H. Plasmodium berghei circumvents immune responses induced by merozoite surface protein 1- and apical membrane antigen 1-based vaccines. PloS one. 2010;5(10):e13727. doi: 10.1371/journal.pone.0013727. PubMed PMID: 21060850; PubMed Central PMCID: PMC2965677.

6. Alaro JR, Partridge A, Miura K, Diouf A, Lopez AM, Angov E, et al. A chimeric Plasmodium falciparum merozoite surface protein vaccine induces high titers of parasite growth inhibitory antibodies. Infection and immunity. 2013;81(10):3843–54. doi: 10.1128/IAI.00522-13. PubMed PMID: 23897613; PubMed Central PMCID: PMC3811772.

7. Burns JM, Jr., Miura K, Sullivan J, Long CA, Barnwell JW. Immunogenicity of a chimeric Plasmodium falciparum merozoite surface protein vaccine in Aotus monkeys. Malar J. 2016;15:159. doi: 10.1186/s12936-016-1226-5. PubMed PMID: 26975721; PubMed Central PMCID: PMC4791798.

8. Sheikh IH, Kaushal DC, Chandra D, Kaushal NA. Immunogenicity of a plasmid DNA vaccine encoding 42kDa fragment of Plasmodium vivax merozoite surface protein-1. Acta tropica. 2016;162:66–74. doi: 10.1016/j.actatropica.2016.06.013. PubMed PMID: 27311385.

9. Neal AT, Jordan SJ, Oliveira AL, Hernandez JN, Branch OH, Rayner JC. Limited variation in vaccine candidate Plasmodium falciparum Merozoite Surface Protein-6 over multiple transmission seasons. Malar J. 2010;9:138. doi: 10.1186/1475-2875-9-138. PubMed PMID: 20497564; PubMed Central PMCID: PMC2881939.

10. Sinden RE, Billingsley PF. Plasmodium invasion of mosquito cells: hawk or dove? Trends in parasitology. 2001;17(5):209–12. Epub 2001/04/27. PubMed PMID: 11323288.

11. Li J, Wang X, Zhang G, Githure J, Yan G, James AA. Genome-block expression-assisted association studies discover malaria resistance genes in *Anopheles gambiae*. Proceedings of the National Academy of Sciences of the United States of America. 2013;110(5):20675–80.

12. Zhang G, Niu G, Franca CM, Dong Y, Wang X, Butler NS, et al. Anopheles midgut FREP1 mediates Plasmodium invasion. The Journal of biological chemistry. 2015;290(27):16490–501. Epub 2015/05/21. doi: 10.1074/jbc.M114.623165. PubMed PMID: 25991725.

13. Niu G, Wang B, Zhang G, King J, Cichewicz RH, Li J. Targeting mosquito FREP1 with a fungal metabolite blocks malaria transmission. Scientific Reports. 2015;5(14694). PubMed PMID: 25991725; PubMed Central PMCID: PMC4505404.

14. Niu G, Franc AC, Zhang G, Roobsoong W, Nguitragool W, Wang X, et al. The fibrinogen-like domain of FREP1 protein is a broad-spectrum malaria transmission-blocking vaccine antigen. The Journal of biological chemistry. 2017;292(28):11960–9. Epub 2017/05/24. doi: 10.1074/jbc.M116.773564. PubMed PMID: 28533429; PubMed Central PMCID: PMCPMC5512087.

15. Niu G, Franca C, Zhang G, Roobsoong W, Nguitragool W, Wang X, et al. The fibrinogen-like domain of FREP1 protein is a broad-spectrum malaria transmission-blocking vaccine antigen. The Journal of biological chemistry. 2017;292(28):11960–9. Epub 2017/05/24. doi: 10.1074/jbc.M116.773564. PubMed PMID: 28533429; PubMed Central PMCID: PMCPMC5512087.

16. Dong Y, Simoes ML, Marois E, Dimopoulos G. CRISPR/Cas9 -mediated gene knockout of Anopheles gambiae FREP1 suppresses malaria parasite infection. PLoS Pathog. 2018;14(3):e1006898. Epub 2018/03/09. doi: 10.1371/journal.ppat.1006898. PubMed PMID: 29518156; PubMed Central PMCID: PMCPMC5843335.

17. Li J, Clinkenbeard KD, Ritchey JW. Bovine CD18 identified as a species specific receptor for Pasteurella haemolytica leukotoxin. Veterinary microbiology. 1999;67(2):91–7. Epub 1999/07/22. PubMed PMID: 10414364.

18. Ghosh AK, Dinglasan RR, Ikadai H, Jacobs-Lorena M. An improved method for the in vitro differentiation of Plasmodium falciparum gametocytes into ookinetes. Malaria journal. 2010;9:194. Epub 2010/07/10. doi: 10.1186/1475-2875-9-194. PubMed PMID: 20615232; PubMed Central PMCID: PMCPMC2909250.

19. Tiong HK, Hartson SD, Muriana PM. Comparison of Surface Proteomes of Adherence Variants of Listeria Monocytogenes Using LC-MS/MS for Identification of Potential Surface Adhesins. Pathogens. 2016;5(2). Epub 2016/05/20. doi: 10.3390/pathogens5020040. PubMed PMID: 27196934; PubMed Central PMCID: PMCPMC4931391.

20. Morrissette NS, Sibley LD. Cytoskeleton of apicomplexan parasites. Microbiol Mol Biol Rev. 2002;66(1):21-38; table of contents. Epub 2002/03/05. PubMed PMID: 11875126; PubMed Central PMCID: PMCPMC120781.

21. Patra KP, Vinetz JM. New ultrastructural analysis of the invasive apparatus of the Plasmodium ookinete. The American journal of tropical medicine and hygiene. 2012;87(3):412–7. doi: 10.4269/ajtmh.2012.11-0609. PubMed PMID: 22802443; PubMed Central PMCID: PMC3435341.

22. Invitrogen. Bac-to-Bac® Baculovirus Expression System 2015. Available from: https://tools.thermofisher.com/content/sfs/manuals/bactobac_man.pdf.

23. QIAGEN. The QIAexpressionist: A handbook for high-level expression and purification of 6XHis-tagged proteins. Hilden, Germany: QIAGEN; 2003.

24. Niu G, Wang B, Zhang G, King J, Cichewicz RH, Li J. Targeting mosquito FREP1 with a fungal metabolite blocks malaria transmission. Scientific Reports. 2015;5(Oct 6, 2015). doi: 10.1038/srep14694.

